# Systematic analyses of factors required for adhesion of *Salmonella enterica* serovar Typhimurium to corn salad (*Valerianella locusta*)

**DOI:** 10.1101/855726

**Authors:** Laura Elpers, Juliane Kretzschmar, Sean-Paul Nuccio, Andreas J. Bäumler, Michael Hensel

## Abstract

*Salmonella enterica* is a foodborne pathogen leading to gastroenteritis and is commonly acquired by consumption of contaminated food of animal origin. However, numbers of outbreaks linked to the consumption of fresh or minimally processed food of non-animal origin are increasing. New infection routes of *S. enterica* by vegetables, fruits, nuts and herbs have to be considered. This leads to special interest in *S. enterica* interactions with leafy products, e.g. salads, that are consumed unprocessed. The attachment of *S. enterica* to salad is a crucial step in contamination, but little is known about the bacterial factors required and mechanisms of adhesion. *S. enterica* possesses a complex set of adhesive structures whose functions are only partly understood. Potentially, *S. enterica* may deploy multiple adhesive strategies for adhering to various salad species, and other vegetables. Here, we systematically analyzed the contribution of the complete adhesiome, of LPS, and of flagella-mediated motility of *S. enterica* serovar Typhimurium (STM) in adhesion to corn salad. We deployed a reductionist, synthetic approach to identify factors involved in the surface binding of STM to leaves of corn salad with particular regard to the expression of all known adhesive structures using the Tet-on system. This work reveals the contribution of Saf fimbriae, type 1 secretion system-secreted BapA, an intact LPS, and flagella-mediated motility of STM in adhesion to corn salad leaves.

Importance

Human gastrointestinal pathogens are often transmitted by animal products, but recent outbreaks show increasing importance of vegetables as source of infection by pathogenic *E. coli* or *Salmonella enterica.* The mechanisms of binding of *S. enterica* to vegetables such as salad are only poorly understood. We established an experimental model system to systematically investigate the role of adhesive structures of *S. enterica* serovar Typhimurium in binding to corn salad leaves. The contributions of all members of the complex adhesiome, flagella, and O-antigen were evaluated. We identified that Saf fimbriae, type 1 secretion system-secreted BapA, an intact LPS, and flagella-mediated motility contribute to adhesion of *Salmonella* to corn salad leaves. These results will enable future investigations on factors contributing to contamination of vegetables under agricultural conditions.

## Introduction

*Salmonella enterica* is one of the main bacterial pathogens leading to foodborne illnesses and thousands of fatal cases worldwide (1). Depending on the serovar, *Salmonella enterica* causes gastroenteritis (non-typhoidal strain, e.g. Typhimurium), or typhoid fever (typhoidal strains, e.g. Typhi and Paratyphi). Focus has historically been on infection routes of *Salmonella* by animal products, although in recent years, an increasing number of infections caused by vegetable fresh produce have been reported. In addition to pathogenic *Escherichia coli* (e.g*. E. coli* O157:H7) or *Listeria monocytogenes*, *Salmonella enterica* is also involved in such plant-associated infections (2–4). Several outbreaks were associated with contaminated vegetables (e.g. tomatoes, salad), fruits (e.g. watermelons, berries), nuts, herbs (e.g. basil), and sprouts (5, 6). Fresh produce can be contaminated either through cultivation (contaminated irrigation water or fertilizer), or during handling and processing. *S. enterica* may adhere to leaves and roots, colonize the plant, and further internalize into the plant tissue. Once inside the plant, *S. enterica* potentially replicate and persist (7, 8). Endophytic colonization by *Salmonella* cannot be removed by surface washing, and bacteria will thus be ingested if food is consumed after minimal processing.

The differences in interaction of *Salmonella* with plants or animals have to be investigated to better understand infection of plant-based products by *S. enterica*. For the analyses of contamination of salads by *S. enterica*, the leafy part is of special interest. Here, the initial binding of *S. enterica* to salad leaves is a key event in the adhesion and further colonization of salad.

Here we employ *S. enterica* serovar Typhimurium (STM) as the main pathogen causing gastroenteritis. STM possesses a large set of adhesive structures, including 12 chaperone-usher (CU) fimbriae, Curli fimbriae assembled by the nucleation-precipitation pathway, two type 1 secretion system (T1SS)-secreted adhesins (BapA and SiiE), and three type 5 secretion system (T5SS)-secreted adhesins (MisL, ShdA and SadA). Further, two outer membrane proteins (OMP) are known with putative adhesive features (PagN and Rck). In addition, motility and chemotaxis mediated by flagellar rotation, as well as the adhesive effect of the lipopolysaccharide (LPS) layer, must be taken into consideration (9, 10). The specific binding properties of only a few adhesive structures of *S. enterica* are known, and thus no educated guess can be made in regard to possible interactions with salad leaves.

Several studies have investigated the adhesion of *S. enterica* serovars to various species of salad (7, 11–18) with focus on individual adhesion factors. These studies revealed the involvement of flagella and motility as well as further virulence-associated genes in adhesion to salad. Further, the impact of different salad species was evaluated. Nevertheless, a major obstacle in these analyses of *S. enterica* adhesion to vegetables was a lack of knowledge for the conditions necessary to express the pathogen’s various adhesins. Indeed, only a minor proportion of adhesins are known to be expressed under laboratory conditions. It can thus be speculated that a subset of adhesins is expressed under environmental conditions outside of a warm-blooded host organism, although a systematic analysis of such expression is pending. To circumvent this limitation and to functionally express the entire adhesiome of STM, we recently devised a simple and robust approach based on the use of the P*_tetA_* promoter and induction by the non-antibiotic tetracycline derivative AHT (19). In the present study, we deploy this technique to investigate the contribution of the various adhesive structures of STM to adhesion to the surface of corn salad leaves.

To our knowledge, we have analyzed the impact of all adhesive structures of STM in adhesion to corn salad. Moreover, we have found factors that are involved in the adhesion of STM to salad. With this knowledge, we are potentially able to devise defensive strategies in growing, harvesting and processing fresh produce in order to decrease the incidence of *Salmonella* infections.

## Results

We deployed a reductionist, synthetic approach to identify factors that contribute to the surface binding of *Salmonella enterica* serovar Typhimurium (STM) to leaves of corn salad. As with all *S. enterica* serovars studied so far, STM possesses a complex adhesiome. As most of these adhesins are not expressed under laboratory conditions, we expressed the various operons or genes encoding adhesins ectopically under control of a tetracycline-inducible promoter, as previously described (19). Strains harboring these adhesin-expressing plasmids were subsequently tested for their contribution to adhesion to corn salad. The infection of corn salad grown under aseptic conditions by STM was performed as described schematically in Figure S 1.

Prior to analyzing the contribution of adhesive structures in adhesion to corn salad, we tested different deletion strains for their suitability as the strain background for subsequent experiments. The laboratory conditions for native expression of only a few adhesins such as *fim* fimbriae are known. Moreover, the expression of a fimbrial adhesin can impact the expression of other systems, including other adhesins (20, 21). To avoid potential interference by these factors, we generated a strain lacking all 12 CU fimbriae (SR11 Δ12). Furthermore, a strain was generated lacking all known and putative adhesive structures in SR11 (Δ*fimAICDHF* Δ*stbABCD* Δ*sthABCDE* Δ*stfACDEFG* Δ*stiABCH* Δ*bcfABCDEFGH* Δ*safABCD* Δ*pefACDorf5orf6* Δ*stcABCD* Δ*stjEDCBA* Δ*stdAB* Δ*lpfABCDE*::KSAC Δ*misL* Δ*sadA* Δ*shdA* ΔSPI4 Δ*bapABCD* Δ*rck* Δ*pagN* Δ*csgBAC-DEFG*) which we termed SR11 Δ20. Under the assay conditions, both SR11 Δ12 and SR11 Δ20 showed the same level of adhesion to corn salad as WT SR11 (Figure 1A). Therefore, we decided to use SR11 Δ12 in all further experiments to avoid any background expression of CU fimbriae during our assays. Furthermore, SR11 Δ12 strains with additional deletions of single adhesive structures showed no altered levels of adhesion compared to SR11 Δ12, except for deletion of *Salmonella* pathogenicity island 4 (SPI4) and of *bapABCD* (Figure 1B). The deletion strain defective in SPI4, lacking SiiE, the corresponding T1SS, as well as accessory proteins, showed increased adhesion (129% on average). The loss of adhesin BapA and its cognate T1SS BapBCD (Δ*bapABCD*) led to significantly decreased adhesion (65% on average). Of interest, BapA was not detected on the bacterial surface in 3.5 h subcultures of parental strain SR11 Δ12 (Figure S 2C-D).

**Figure 1:**
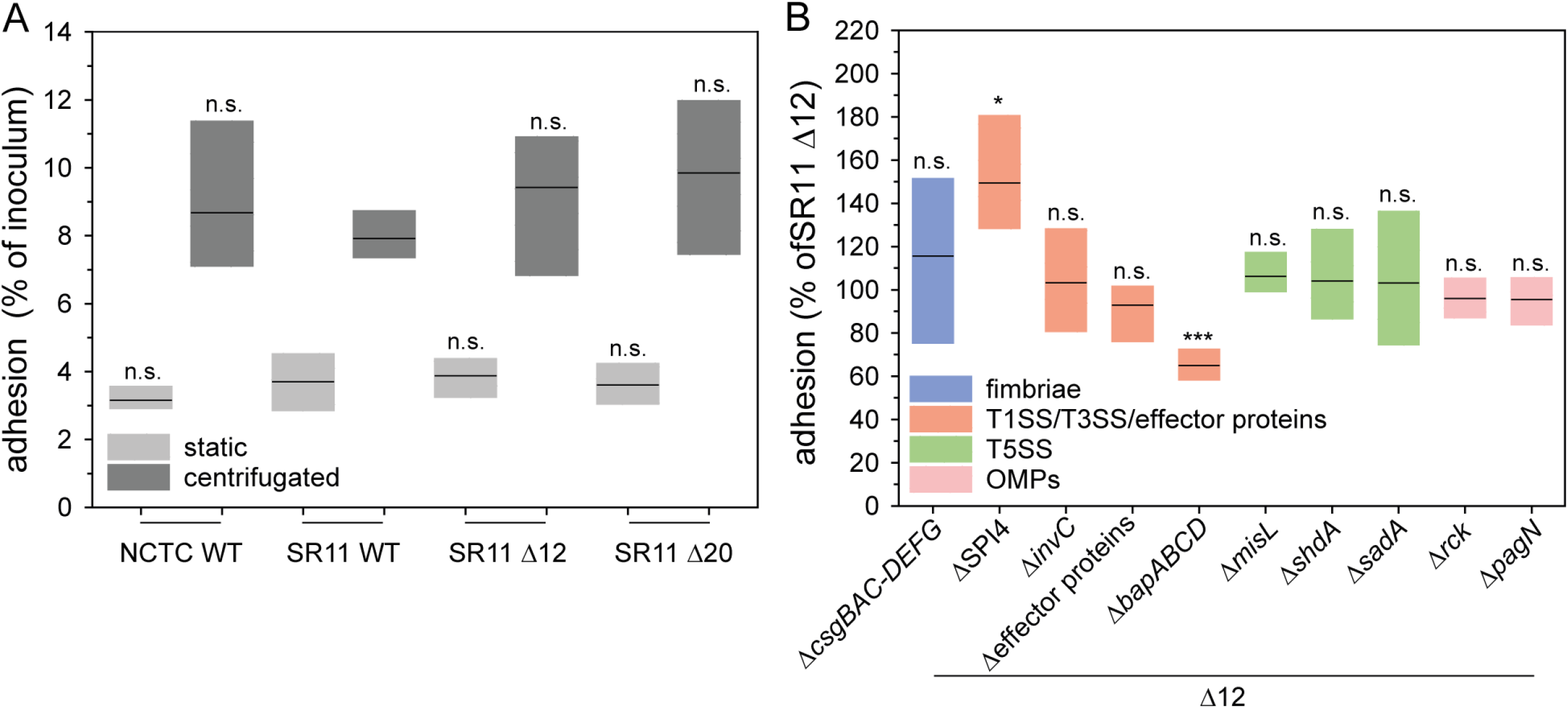
Comparison of *Salmonella* NCTC WT, SR11 WT, SR11 Δ12 and SR11 Δ20 and impact of deficits in genes encoding for putative adhesive structures and effector proteins of SPI1-T3SS STM adhesion to corn salad. Corn salad grown under aseptic conditions was infected with STM strain NCTC WT, SR11 WT, SR11 Δ12 and SR11 Δ20 (A) and with SR11 Δ12 with various deletions in genes encoding putative adhesive structures and effector proteins of SPI1-T3SS (Δ*sopA* Δ*sopB* Δ*sopD* Δ*sopE2* Δ*sipA* = Δeffector proteins) (B). Bacteria were subcultured for 3.5 h (1:31) and diluted in PBS for infection of corn salad. After infection for 1 h corn salad segments were washed three times to remove non-adherent bacteria. For the quantification of adherent bacteria, corn salad leaf discs were homogenized in PBS containing 1% deoxycholate, and serial dilutions of homogenates and inoculum were plated onto MH agar plates for the quantification of CFU. Adhesion rates in % of inoculum were determined by the ratio of CFU in inoculum and homogenate, and adherent bacteria normalized to SR11 Δ12 set as 100% adhesion. Shown are the distributions of three biological replicates represented as box plots with median. Statistical significances were calculated with Student’s *t*-test and are indicated as follows: n.s., not significant; *, *p* ≤ 0.05; **, *p* < 0.01; ***, *p* < 0.001.

### Contribution of fimbrial adhesins to adhesion to corn salad

For most of the 12 CU fimbriae, little is known about their native expression and binding properties (22). All operons encoding CU fimbriae consist of at least a fimbrial main subunit, a specific periplasmic chaperone, and a specific usher located in the outer membrane (23). The most prominent and best-studied fimbriae are type 1 fimbriae encoded by the *fim* operon (*fimAICDHF*). Type 1 fimbriae are natively expressed under static conditions and mediate binding to mannosylated proteins (24).

We analysed adhesion to corn salad after P*_tetA_*-induced expression of various CU fimbriae (Figure 2A). The assay revealed distinct phenotypes of binding to corn salad. Expression of certain CU fimbriae by STM (Lpf, Bcf, Sth, Std and Stj) resulted in adhesion levels similar to background strain SR11 Δ12, indicating that these adhesins do not have cognate ligands on corn salad. Adhesion to corn salad was impaired after expression of Fim, Pef, Stc and Stb fimbriae (53%; 72%; 58% and 60% mean adhesion rates, respectively, compared to SR11 Δ12), while expression of Sti fimbriae resulted in slightly, but not significantly decreased adhesion. By contrast, AHT-induced expression of Saf and Stf fimbriae led to increased adhesion (166% and 116% mean adhesion rates, respectively). A clear contribution of Saf fimbriae in adhesion to corn salad was confirmed by the non-induced control, exhibiting no altered adhesion level compared to background strain SR11 Δ12. Of note, a non-significant increase in adhesion was observed for Stf fimbriae in the absence of the inducer AHT, which was comparable to the AHT-induced samples. As such, a clear role for Stf fimbriae in STM’s adhesion to corn salad cannot be ascribed.

**Figure 2:**
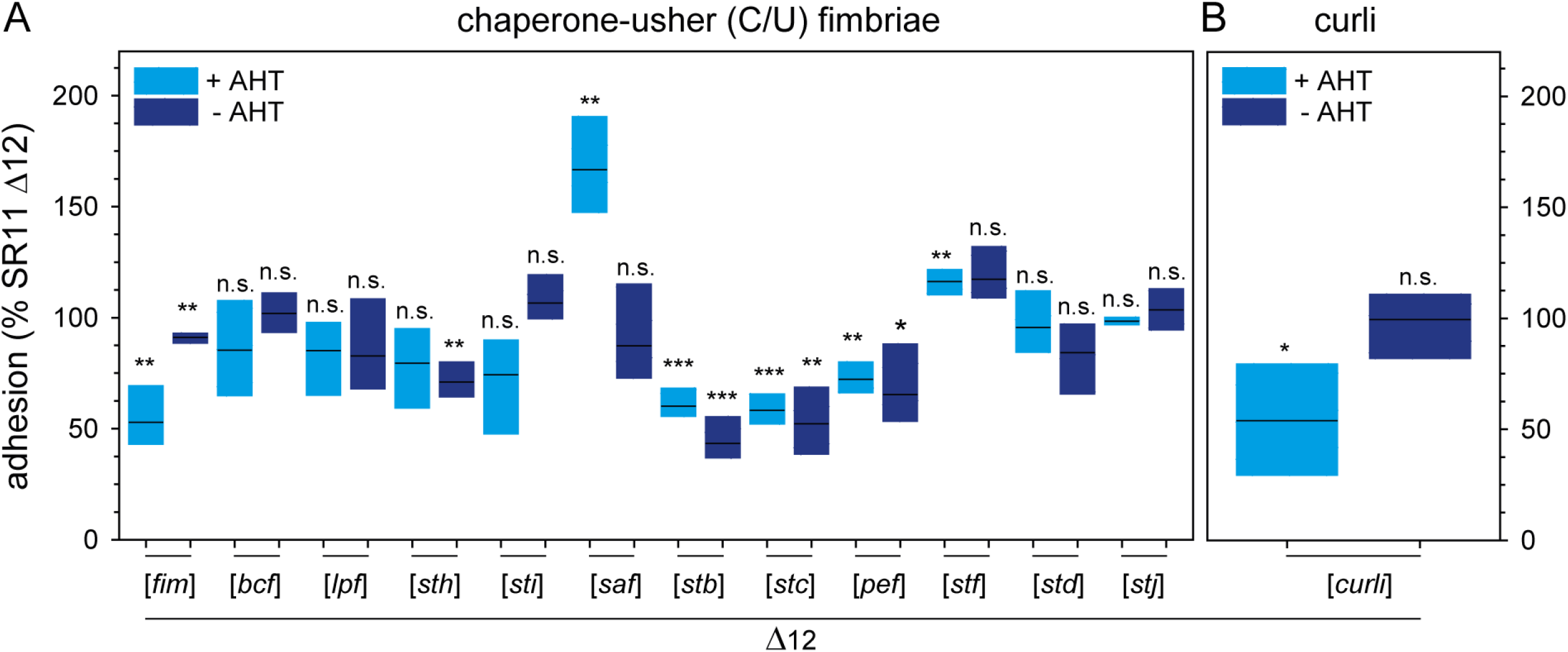
Impact of chaperone-usher fimbriae and Curli fimbriae expression on STM adhesion to corn salad. Sterile grown corn salad was infected with *S. enterica* serovar Typhimurium strain SR11 Δ12 with the expression of various chaperone-usher fimbriae (A) and the expression of Curli fimbriae in the according deletion mutant (B). Expression of fimbriae were induced by 10 ng/ml AHT for 3.5 h in subculture. The adhesion and statistical significances were determined as described in Figure 1.

Curli fimbriae are known to be involved in biofilm formation (25) and are encoded by two divergent operons, *csgBAC* and *csgDEFG*, with assembly occurring via the nucleation-precipitation pathway. AHT-induced expression of Curli fimbriae showed a decreased adhesion to corn salad, whereas without AHT induction no altered adhesion was observed (Figure 2B).

### Contribution of T1SS-secreted non-fimbrial adhesins to adhesion to corn salad

The SPI4 locus (*siiABCDEF*, *<u>S</u>almonella <u>i</u>ntestinal <u>i</u>nfection*) encodes the giant adhesin SiiE which is secreted to the bacterial surface by the T1SS SiiCDF (26). The two accessory proteins SiiA and SiiB are located in the inner membrane and presumably function as a proton-conductive channel (27). SiiE is known as the largest protein in STM, with 53 repetitive bacterial Ig domains (BIg) and a molecular mass of 595 kDa. Moreover, SiiE exhibits binding specificity for glycostructures with terminal N-acetyl-glucosamine (GlcNAc) and 2,3-linked sialic acid (28). SiiE mediates the first contact of *Salmonella* to polarized epithelial cells of mammalian hosts (e.g. MDCK cells), enabling subsequent invasion mediated by the SPI1-encoded T3SS and various effector proteins (29, 30). As generation of a vector for Tet-on expression of the *sii* operon turned out to be problematic, we deployed an alternative approach to control expression of the native *sii* operon. Enhanced surface expression of SiiE was achieved by AHT-induced overexpression of *hilD*, the central transcriptional activator of the SPI1/SPI4 regulon (31). We observed that increased amounts of SiiE on the bacterial surface led to decreased adhesion to corn salad (77% mean, Figure 3A) compared to SR11 Δ12 with native expression of SiiE in 3.5 h subcultures (Figure S 2AB). Without induction by AHT and therefore with almost native SiiE expression, no differences in adhesion compared to the background strain SR11 Δ12 was observed. Since the expression of the regulator *hilD* also influences the expression of the SPI1-encoded T3SS and its effector proteins, the plasmid encoding *hilD* was tested under control of the Tet-on system in further SPI1 and SPI4 deletion mutants. Overexpression of *hilD* in a SPI4 deletion mutant led to a significantly decreased adhesion of 53% in average, indicating that the SPI1-T3SS rather than SiiE itself interferes with the adhesion to corn salad. This was further confirmed by an increased adhesion rate of a strain lacking *invC* (ATPase subunit of SPI1-T3SS), and thereby the SPI1-T3SS, harboring a plasmid for *hilD* overexpression (153%). The deletion of *invC* alone, as well as deletion of the effector proteins SopA, SopB, SopD, SopE2 and SipA (= Δ5; Figure 1B) did not alter adhesion, leading to the hypothesis that the SPI1-T3SS affects adhesion to corn salad.

**Figure 3:**
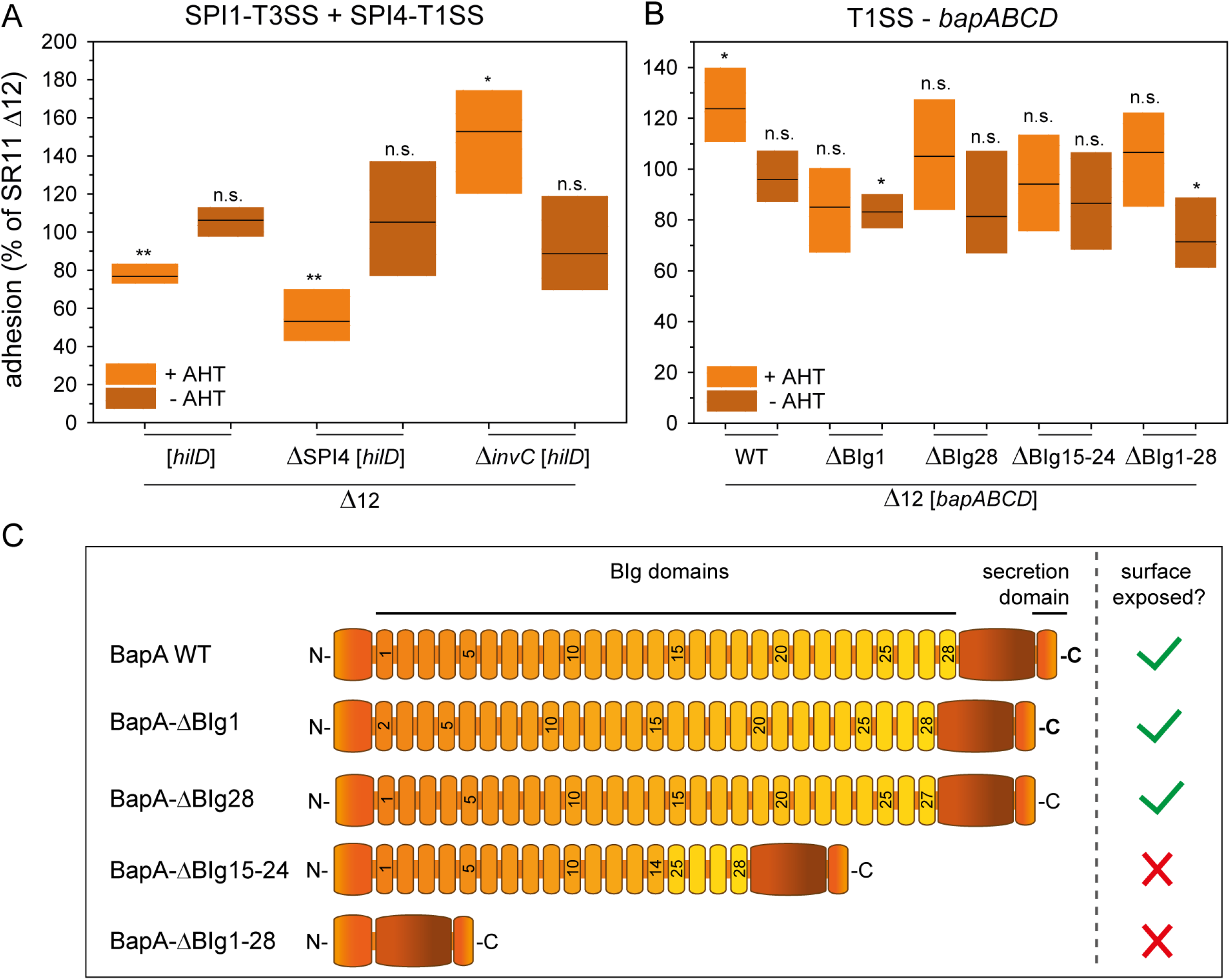
Impact of T1SS-secreted adhesins and *hilD* expression on STM adhesion to corn salad. Corn salad grown under aseptic conditions was infected with STM strain SR11 Δ12 with the overexpression of the regulator *hilD* for the analysis of the SPI4-encoded, T1SS-secreted adhesin SiiE and the SPI1-encoded T3SS (A). In addition, SR11 Δ12 Δ*bapABCD* strains expressing AHT-induced, T1SS-secreted wild-type adhesin BapA, or the indicated BapA truncation mutants were tested (B). The adhesion and the statistical significances were determined as described in Figure 1. (C) gives schematic overview of truncated BapA forms used in adhesion assays.

The *bap* operon (*bapABCD*, *<u>b</u>iofilm-<u>a</u>ssociated <u>p</u>rotein*) encodes for a T1SS including BapB (outer membrane protein), BapC (ATPase) and BapD (membrane fusion protein) which is necessary for the secretion of the adhesin BapA to the bacterial surface. The T1SS-secreted adhesin BapA has a molecular mass of 386 kDa, contains 28 BIg domains, and is involved in biofilm formation (32). AHT-induced expression of the *bap* operon led to increased adhesion to corn salad (124% mean, Figure 3B), whereas no significant differences were observed without AHT induction.

To gain further insight into which structural features of BapA are essential for adhesion, we generated plasmids for Tet-on expression of *bapABCD* that encode BapA with deletions of BIg domains of various extent. Synthesis and secretion of truncated forms of BapA were confirmed by flow cytometry (Figure S 2C-D), and indicated that deletion of BIg1-28 and BIg15-24 ablated the surface expression of BapA. This observation is in line with the adhesion assay results for strains expressing BapA harboring a deletion of BIg1-28 or BIg15-24, which showed no increased adhesion to corn salad. Thus, the loss of BapA surface expression resulted in adhesion levels comparable to SR11 Δ12. By contrast, truncated forms of BapA with deletion of only one BIg domain, either BIg1 or BIg28, were detected on the bacterial surface by flow cytometry. Moreover, in adhesion assays, no increased adhesion was observed compared to wild type BapA. Hence, the BIg1 and BIg28 domains might be relevant for proper binding to corn salad by BapA.

### Contribution of autotransported adhesins to adhesion to corn salad

STM expresses three autotransported adhesins: MisL, ShdA and SadA. MisL and ShdA are monomeric adhesins, whereas SadA belongs to the class of trimeric adhesins. Previous studies have shown that MisL and ShdA are involved in binding to fibronectin, with impacts intestinal infection of mice (33, 34). SadA is possibly involved in adhesion to CaCo2 cells, as well as in biofilm formation, but only in a strain background with altered LPS structure (35). AHT-induced expression of *misL* did not alter adhesion to corn salad (Figure 4A). In contrast, the AHT-induced expression of *shdA* led to a decreased average adhesion of 67%, whereas the non-induced strain displayed no changes in adhesion. The AHT-induced expression of *sadA* and its chaperone *sadB* led to a slight, but non-significantly decreased adhesion (79% mean). Although we observed significantly higher adhesion (158% mean) without AHT induction, SadA surface expression was not detected by flow cytometry in non-induced samples (Figure S 2E-F).

**Figure 4.**
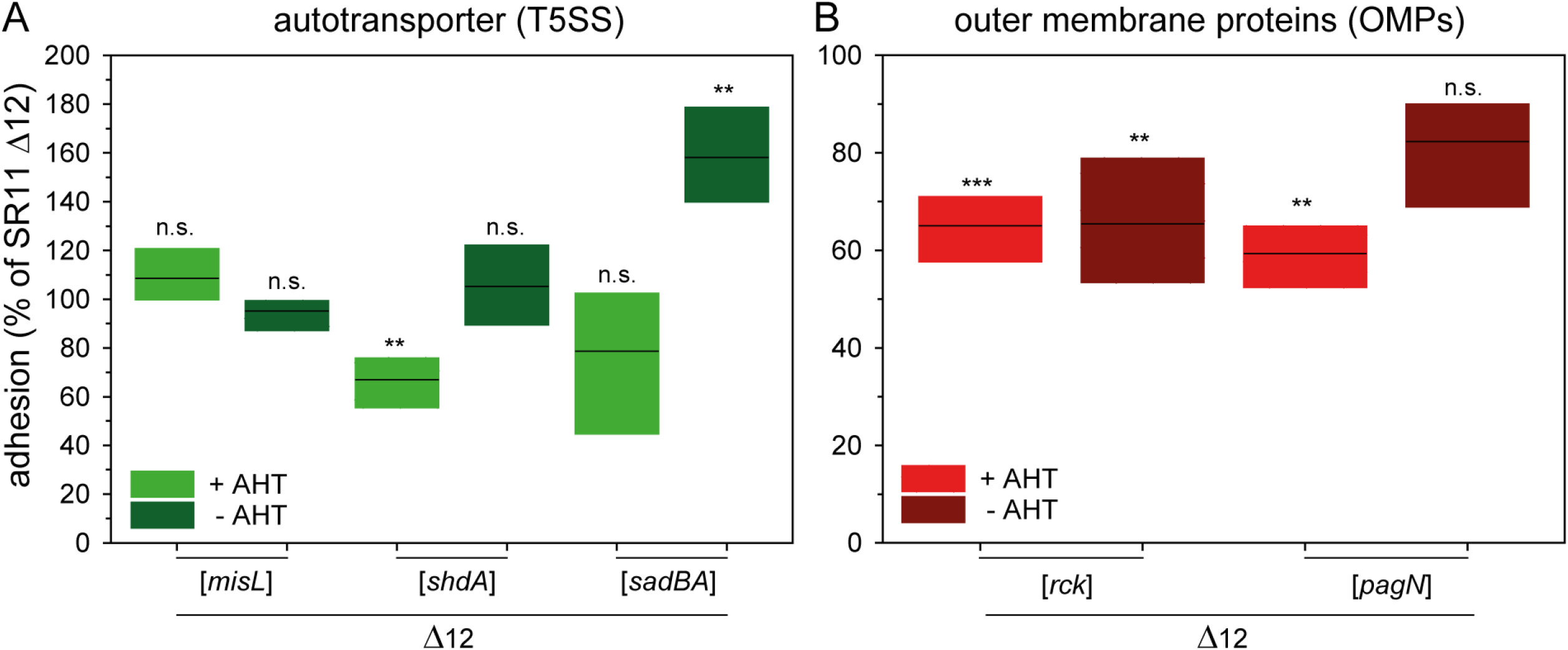
Impact of T5SS-secreted adhesins and of outer membrane proteins on STM adhesion to corn salad. Corn salad grown under aseptic conditions was infected with STM strain SR11 Δ12 expressing the different T5SS-secreted adhesins MisL, ShdA and SadA induced by AHT in the according deletion mutants Δ*misL*, Δ*shdA* and Δ*sadA* (A). For the analysis of outer membrane proteins SR11 Δ12 expressing *rck* and *pagN* by induction of AHT was used in the respective deletion mutants Δ*rck* and Δ*pagN* (B). The adhesion and the statistical significances were determined as described in Figure 1.

### Contribution of OMP adhesins to adhesion to corn salad

The OMPs Rck and PagN are adhesive structures, and an involvement in SPI1-T3SS-independent invasion of epithelial cells has been reported (36, 37). AHT-induced expression of *rck* led to a significantly decreased adhesion to corn salad (65% mean), although even the non-induced sample exhibited decreased adhesion (65% mean, Figure 4B). In a previous study, Western blot analyses confirmed absence of expression of Rck in non-induced cultures (19). The AHT-induced expression of PagN exhibited significantly reduced adhesion (59% in average), whereas the non-induced samples showed no altered adhesion level.

### Contribution of flagellar filaments and motility to adhesion to corn salad

The effect of flagella and motility on infection of various plants has been previously investigated for *Salmonella* and other pathogenic bacteria (15, 38, 39). Here we demonstrate the binding properties and the contribution of motility in adhesion to corn salad using four distinct deletion strains. The deletion of *fliC* and *fljB*, resulting in the loss of the flagellar filament, showed a decreased adhesion (50% mean) which could not be restored to background strain level by centrifugation (Figure 5A). This effect may thus be due to an adhesive feature of the flagellar filament, or due to flagella-mediated motility promoting contact to corn salad surfaces. To dissect the contribution of flagella, a *motAB* mutant strain was employed; these strains still produce a flagellar filament, but are unable to energize the flagellar motor and are thus non-motile. The Δ*motAB* strain showed decreased adhesion for static and centrifuged samples (67% and 73% mean). Thus, presence of flagella without motility does not enable *Salmonella* to bind to corn salad. To gain further insight into how motility contributes to adhesion to corn salad, we deployed mutant strains with defective *cheY*, resulting in a strong bias towards smooth swimming, or defective *cheZ*, resulting in a strong bias for tumbling (Figure 5C). The Δ*cheY* strain showed a decreased adhesion (71% mean) after centrifugation, whereas the deletion of *cheZ* led to a decreased adhesion which is non-significant in static and centrifuged samples. We conclude that proper flagella-mediated motility contributes to adhesion to corn salad surfaces, and this effect is not caused by sole interaction of the flagella filament with the leave surface.

**Figure 5:**
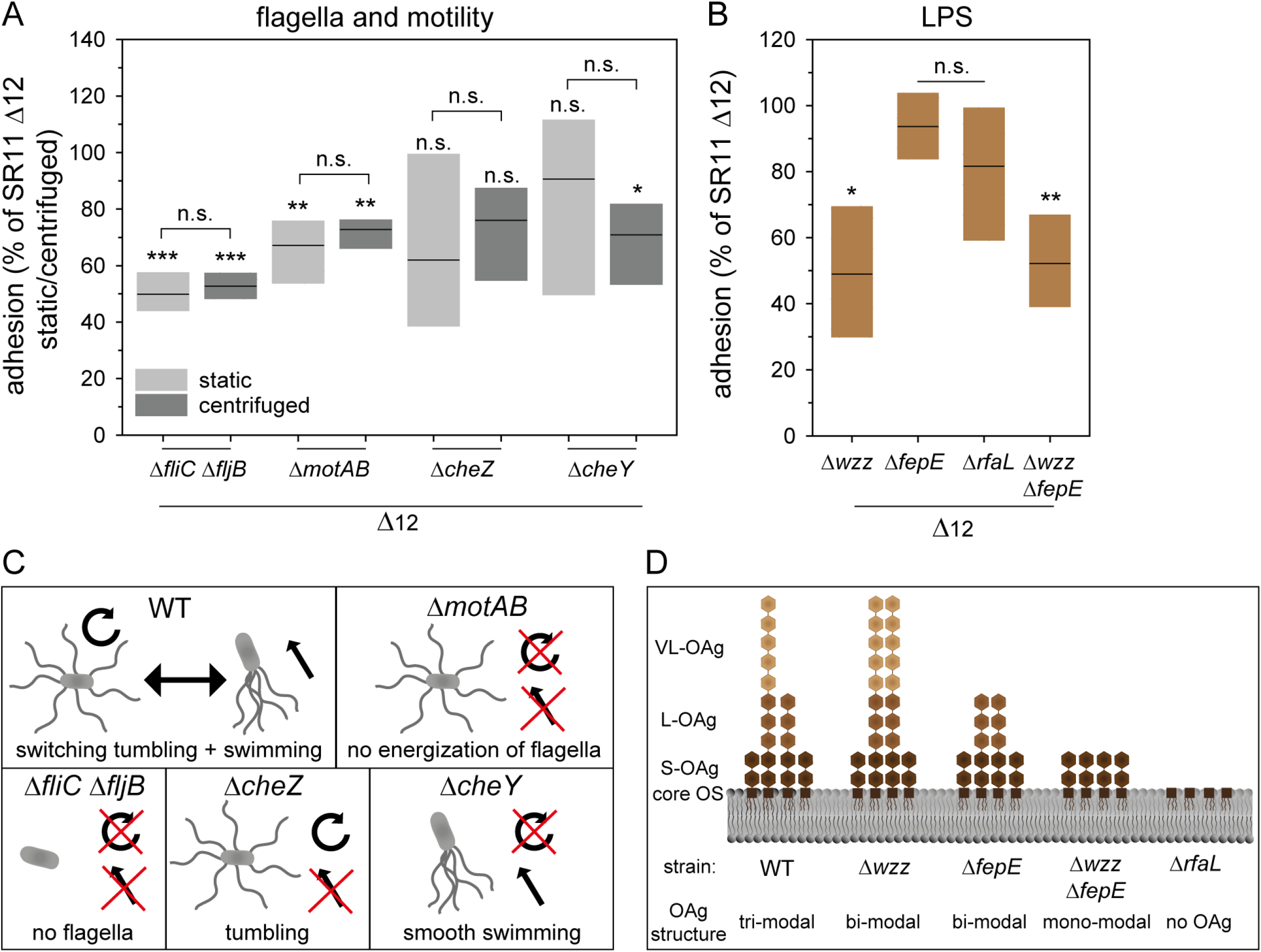
Impact of defect in motility and flagella assembly and deletion of LPS structure on STM adhesion to corn salad. Corn salad grown under aseptic conditions was infected with STM strain SR11 Δ12 with deletion of various motility and flagella-associated genes (A) and deletion of LPS structure-related genes (B). The infection took place either under static conditions or after centrifugation at 500 x g for 5 min to compensate effects of mutations in motility genes. For deletion of genes involved in O-antigen biosynthesis, only static samples are shown here. The adhesion and the statistical significances were determined as described in Figure 1. Models of the resulting phenotype depending on the different deletions in motility flagella assembly and LPS structure are depicted in C) and D). Panel D is based on (44).

### Contribution of O-antigen to adhesion to corn salad

The major constituent of the Gram-negative cell surface is LPS. In addition to stabilization of the cell envelope and protection against various environmental factors, LPS increases the negative charge of the cell envelope and a putative adhesive role has been reported (40). To analyze the impact of LPS in adhesion to corn salad, we used mutant strains lacking various genes involved in the biosynthesis of the O-antigen of LPS. WT *Salmonella* displays a heterogeneous distribution of long chain O-antigens (L-OAg), and very long chain O-antigens (VL-OAg). Deletion of *wzz* results in the homogenous distribution of VL-OAg, deletion of *fepE* in homogenous distribution of L-OAg, and a strain lacking both genes (*wzz fepE*) can only synthesize short O-antigen (S-OAg) (Figure 5D). The deletion of *rfaL* leads to lack of O-antigen, resulting in LPS restricted to the core oligosaccharides.

In this study the deletion of *wzz* and *wzz fepE* led to a decreased adhesion in static (49% and 52%) samples (Figure 5B). The deletion of *rfaL* displayed a decreased adhesion (82% mean) which was non-significant. The strain lacking *fepE* showed no altered adhesion. These data suggest that presence of only VL-OAg, or only S-OAg impairs binding to corn salad, and as a consequence the L-OAg has to be present. The observation that the Δ*rfaL* lacking O-antigen and further led to no significant decrease in the adhesion rate could be explained by binding of the core oligosaccharide to corn salad. All results obtained in this study are summarized in a schematic overview in Figure 6.

**Figure 6:**
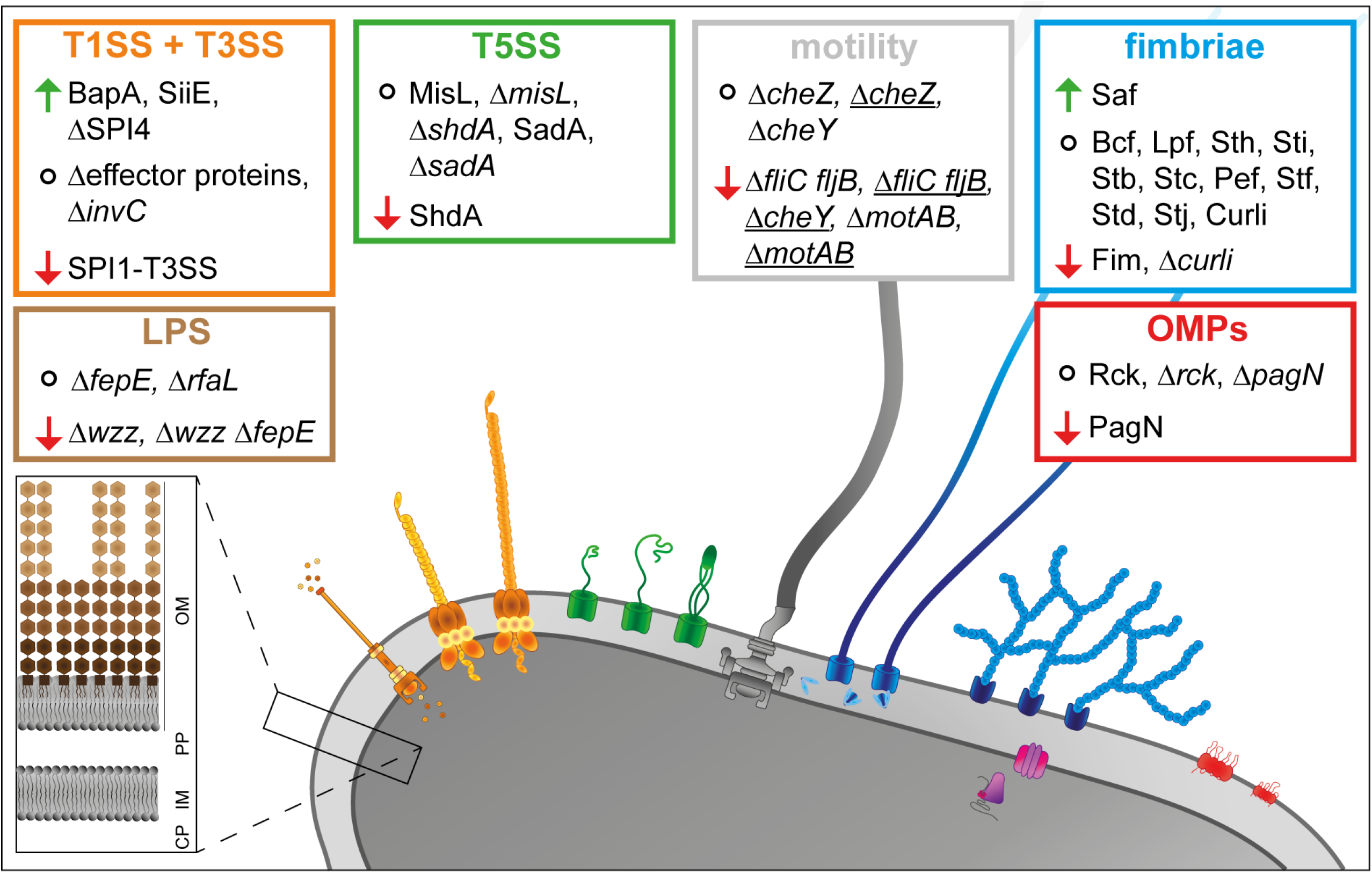
Overview of the impact of the analyzed factors of *Salmonella* Typhimurium in adhesion to corn salad. Static samples, <u>centrifuged samples</u>, ↑ increased adhesion; ↓decreased adhesion; ○ no altered adhesion.

## Discussion

Previous studies investigated adhesion of *Salmonella* Typhimurium to various plants (including leaves, roots and fruits), thereby clarifying which factors are involved in the first step of colonization (11, 38). These studies succeeded in clarifying the first steps of colonization using WT strains or mutant strains defective in single adhesion factors. In addition, most studies on plant-pathogen interactions only tested differences in adhesion of one *Salmonella* isolate to various plant species, or adhesion of various *Salmonella* serovars to one plant species (12, 13, 41). Currently, the question which factors of *Salmonella* are involved in adhesion to plants remains unanswered. As the expression conditions of most adhesive structures are not known, we propose an approach using the Tet-on system to control expression of all putative adhesive structures of STM, one at a time. In this study, the adhesins were tested for their impact in adhesion to corn salad leaves, using a reductionist, synthetic approach to identify factors that possibly contribute to the surface binding of corn salad leaves.

We found that expression of certain adhesins affected adhesion to corn salad leaves, namely Saf fimbriae, T1SS-secreted BapA and T1SS-secreted SiiE. Expression of other adhesive structures, like type 1 fimbriae, T5SS-secreted ShdA and OMP PagN, caused a decreased adhesion, possibly due to occlusion of other structures involved in adhesion. In addition, the deletion of various basic structural features of bacteria, like O-antigen of LPS, or motility by flagella rotation, had impact on the adhesion to corn salad.

Several previous studies showed that absence of the flagella filament had an influence on adhesion to various plants. Whereas C. N. Berger et al. (11) reported a decreased adhesion to basil leaves for a Δ*fliC* Δ*fljB* strain of *Salmonella* Senftenberg, A. L. Iniguez et al. (42) revealed an enhanced colonization of *Arabidopsis* roots for Δ*fliC* Δ*fljB* mutant of STM. Thus, there has to be a clear difference in the role of flagella between colonization of rhizosphere and phyllosphere. For the colonization of roots, the presence of a flagella is apparently obstructive, due to PAMP-triggered immunity by receptor kinase FLS2 recognition of flg22 by (42, 43). For the first contact of *Salmonella* and other pathogenic bacteria to leaf surfaces, the presence of flagella is of crucial importance. To investigate the possible binding of flagella filament, Y. Rossez et al. (39) purified the flagella filament of pathogenic EHEC *E. coli* O157:H7 Sakai, EPEC *E. coli* O127:H6 and non-pathogenic *E. coli* K12 with flagella serotype H48. They showed that the binding of purified flagella filaments to multiple plant lipid species (SQDG (sulphated glycolipid), phosphatidylcholine, phosphatidylglycerol, phosphatidylinositol and phosphatidylethanolamine) results in the assumption of an ionic adhesion by binding to sulphated and phosphorylated plant plasma membrane lipids with negative charge. In addition, *E. coli* strain TUV93-0 Δ*fliC* showed a decreased adhesion to *Arabidopsis* leaves which could be complemented by all three flagella serotypes (39). Possibly, the ionic adhesion of flagella filament represents a conserved mechanism for adhesion to plant leaves among gram-negative bacteria. Despite analyses of flagella filament involvement in adhesion to various plant organs, less is known about the impact of motility. Y. Kroupitski et al. (15) showed that deletion of Δ*cheY* in STM had no consequences for attachment to iceberg lettuce leaves, whereas the internalization of STM was affected. The authors hypothesize that STM cannot reach stomata due to the lack of directed motility. Directed motility conceivably enables STM to sense sucrose near stomata facilitating internalization. Thus, internalization was impaired during an experiment performed in dark with fusicoccin-treated leaves leading to constitutively opened stomata without producing sucrose by photosynthesis (15). In this study, we detected decreased adhesion levels for strains lacking flagella filaments (Δ*fliC* Δ*fljB*), or the energization of flagella rotation (Δ*motAB)* either under conditions of natural contact, or forced contact. We therefore conclude that not only flagella filaments are needed for adhesion to corn salad leaves, but also motility. We observed only moderate effects in adhesion to corn salad leaves in either the absence of CW or CCW rotation, leading to the assumption that the flagella filament and energization of at least CW or CCW rotation is necessary for binding to corn salad leaves.

However, bacteria might utilize directed motility for accumulation near stomata and/or colonization of plant leaves.

The LPS layer of STM and other pathogenic bacteria was often examined with focus on adhesion to, and invasion of mammalian cells, and for the impact in inflammatory responses. The impact of LPS on adhesion to plant leaves, roots and fruits remained unclear. Mutant strains of STM lacking very long O-Ag, or long and very long O-Ag revealed higher levels of invasion of HeLa and MDCK cells, whereas deletion of the whole O-Ag even led to a highly increased adhesion to both cell lines. Despite this virulence advantage for STM, immune escape was reduced due to higher effector protein translocation (44). In contrast to an enhanced adhesion to mammalian cells by an altered LPS structure of STM, we found that an altered LPS structure resulted in decreased adhesion to corn salad leaves. Our findings are in line with a study by H. Jang and K. R. Matthews (45) revealing that a truncated O-Ag in pathogenic *E. coli* O157:H7 decreases the ability to survive and persist on *Arabidopsis* plants as well as on romaine lettuce. In addition to pathogenic bacteria, an intact LPS structure is also important in non-pathogenic bacteria like *Herbaspirillum seropedicae* acting as a symbiont for many agriculturally important plants. An altered LPS structure in *H. seropedicae* led to decreased attachment to maize root surface and to further endophytic colonization (46). These results were also observed for WT *H. seropedicae* when LPS, N-acetyl glucosamine or glucosamine were added to act as competitors for binding sites. Here we showed for the first time the importance of STM LPS in adhesion to leaf surfaces.

Regardless of LPS and motility of STM, adhesion was increased by expression of different adhesins. Saf fimbriae (*<u>S</u>almonella* <u>a</u>typical <u>f</u>imbriae) were the only fimbriae of the CU pathway found in this study to be involved in enhancing STM adhesion to corn salad leaves. O. Salih et al. (47) revealed by electron microscopy the highly flexible linear structure of Saf fimbriae belonging to FG-loop Long fimbriae (FGL). In contrast to rigid, rod-shaped FG-loop Short (FGS) fimbriae exhibiting various subunits with a distal adhesive tip, FG-loop Long fimbriae often only displays two subunits (48). Thereby, the adhesive unit is likely formed by the most numerous subunits. Thus FGL fimbriae, like Saf fimbriae, might bind to a high number of receptors or ligands (47). Nevertheless, binding properties of Saf fimbriae are unknown. Until now, Saf fimbriae were reported to be involved in biofilm formation and in binding to porcine intestine cells IPEC-J2 cells (49). In addition, expression of Saf fimbriae were only observed during infection of murine spleen (50). Genes of the *saf* operon are often pseudogenes in host-restricted *Salmonella* serovars (Typhi, Paratyphi and Gallinarum) (10), indicating their potential contribution in STM in dispersal by farm animals and newly investigated environmental routes, e.g. leafy plants and other vegetables. To gain further insight in contribution of Saf fimbriae adhesion of STM to plants, binding properties of Saf fimbriae have to be investigated, for example by glycan arrays (39, 51), or by a detailed mutagenesis of potential binding domains.

In this study, we showed that both T1SS-secreted adhesins SiiE and BapA contribute to adhesion to corn salad leaves. While SiiE involvement in adhesion to mammalian polarized epithelial cells by binding GlcNAc and sialic acid is well understood (28, 29), a potential role of SiiE in adhesion to plant surfaces is less likely. The tight control of expression of the SPI1/SPI4 regulon by host cell factors would exclude surface expression of SiiE under environmental conditions. A contribution to adhesion was shown for T1SS-secreted adhesin BapA, and BapA contributes to biofilm formation (53), especially for formation of pellicles on air-liquid interphase (32). Furthermore, deletion of BapA led to a decreased mortality in mice infection. Our data obtained after 1 h of infection excluded the possibility of biofilm formation by BapA expressing STM on corn salad leaves. However, specific binding properties are unknown. To gain further insight in binding properties of BapA to corn salad leaves, various truncated forms of BapA were tested. Truncated forms of BapA lacking one BIg domain were surface expressed and showed no autoaggregation. Deletion of one or more BIg domains reduced BapA-dependent adhesion. Thus, we propose a diminished adhesion to corn salad leaves by a shortened BapA. This hypothesis is further supported by the fact that deletion of BIg1, possibly never reaching out of O-Ag layer in WT BapA, results in a similar phenotype as deletion of BIg28, possibly reaching out of LPS layer first in WT BapA. Further characterization of BapA binding to corn salad leaves is necessary thereby investigating importance of proper folding of BapA in presence of Ca^2+^ (54), and specific binding properties. This study showed for the first time that adhesion of STM to corn salad leaves depends on an intact LPS layer, and on flagella-mediated motility. Further, we revealed the involvement in adhesion to corn salad leaves by expression of CU pathway-assembled Saf fimbriae, T1SS-secreted SiiE and T1SS-secreted BapA. To gain further insight in adhesion of STM to salad, additional salad species should be investigated to access if the detected contributing structures are also involved in adhesion to other salad species, or even to leafy plants in general. Moreover, a transcriptomic and proteomic analysis of the involved adhesins could further elucidate environmental conditions or conditions during colonization of plants. We used a synthetic system with controlled expression of one adhesive factor at a time. If the adhesive factors determined here are also expressed and functional under conditions of natural contamination of plants, has to be investigated by further studies.

In summary, this work contributed to identification of STM adhesive factors required for adhesion to plants. To take these studies to a global context and the pathogen-plant interaction under field-like conditions now a more complex experimental setting is needed.

## Materials and Methods

### Bacterial strains and culture conditions

Bacterial strains used in this study are listed in Table 1. Unless otherwise mentioned, bacteria were routinely grown aerobically in LB (lysogeny broth) medium or on LB agar containing antibiotics if required for selection of specific markers. Carbenicillin (Carb), nalidixic acid (Nal), or kanamycin (Km) were used to a final concentration of 50 µg/ml if required for the selection of phenotypes or maintenance of plasmids. Chloramphenicol (Cm) was used at 30 µg/ml. When needed for cloning purposes, X-Gal was added to LB agar at 20 µg/ml. For the induction of the Tet-on system, anhydrotetracycline (AHT) was used at final concentrations of 10 ng/ml or 100 ng/ml.

**Table 1.**
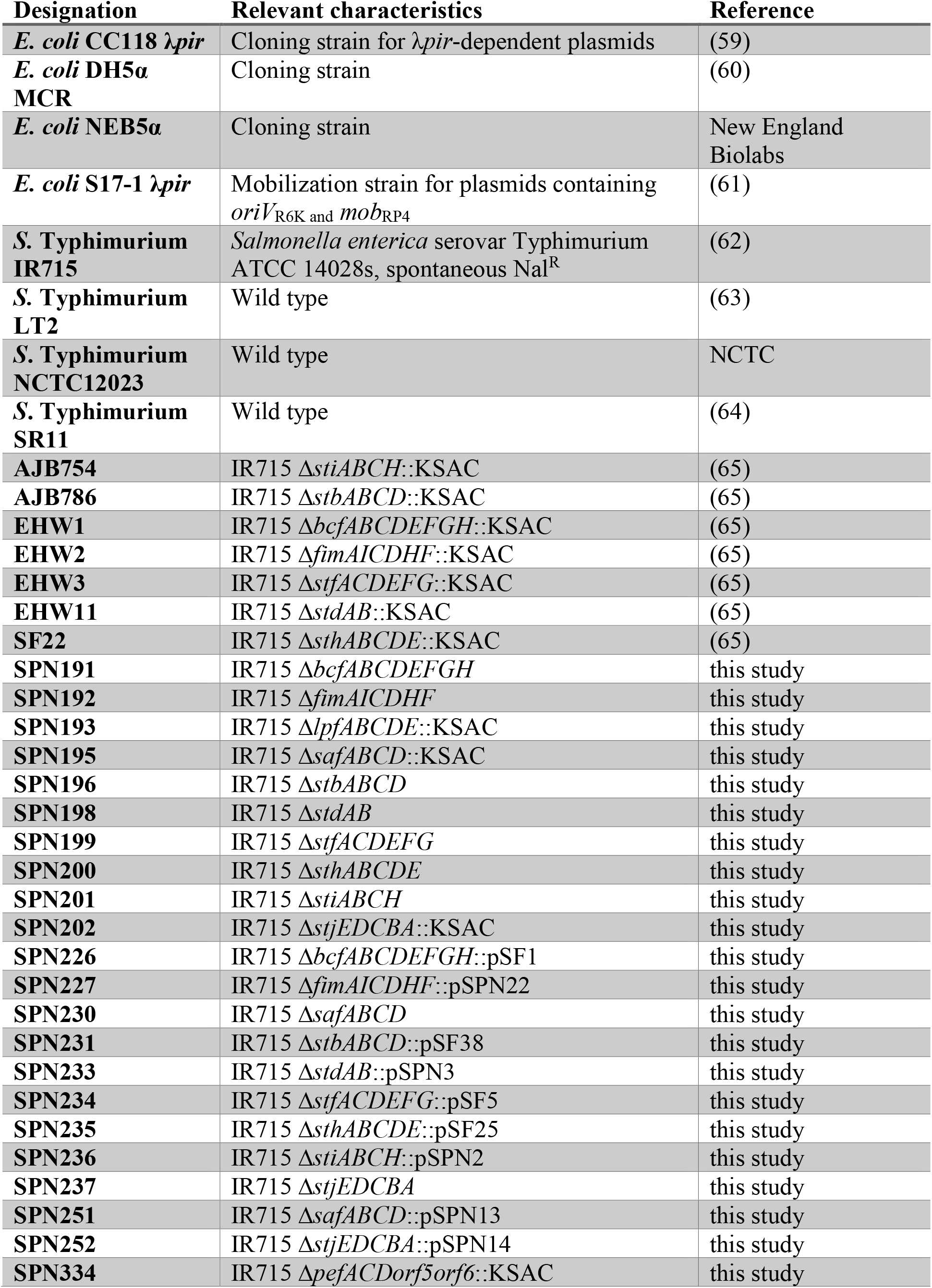
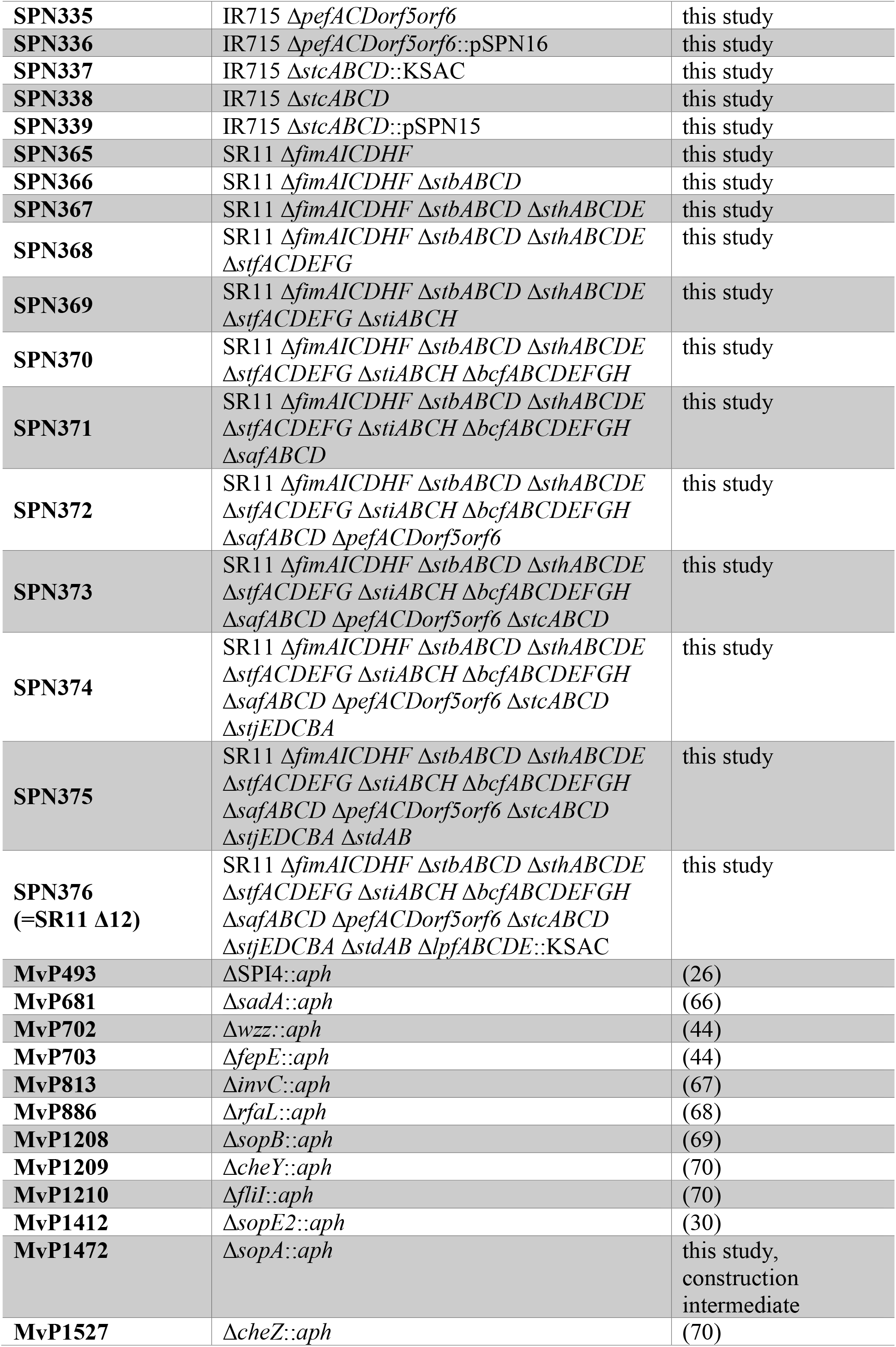
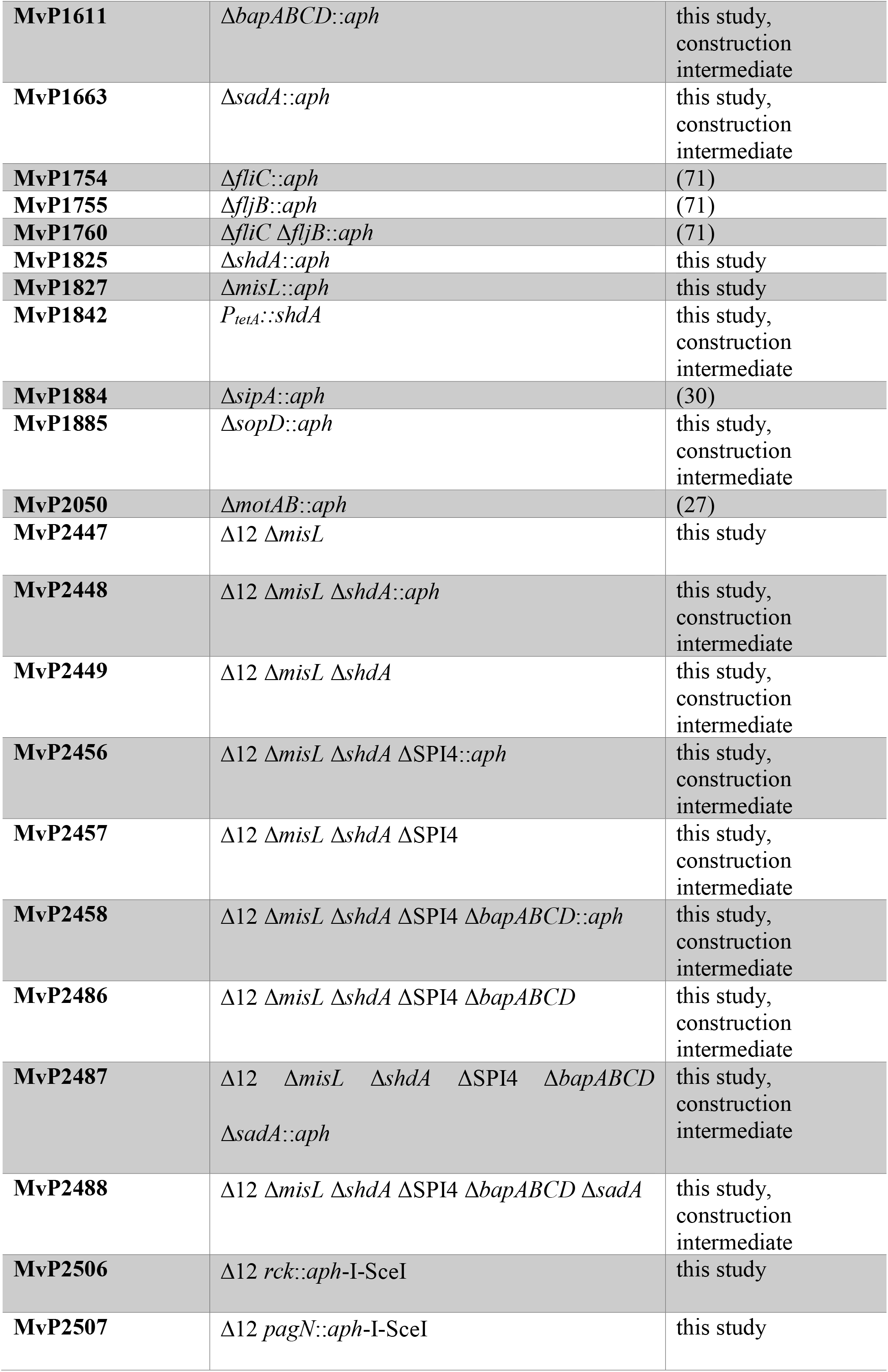
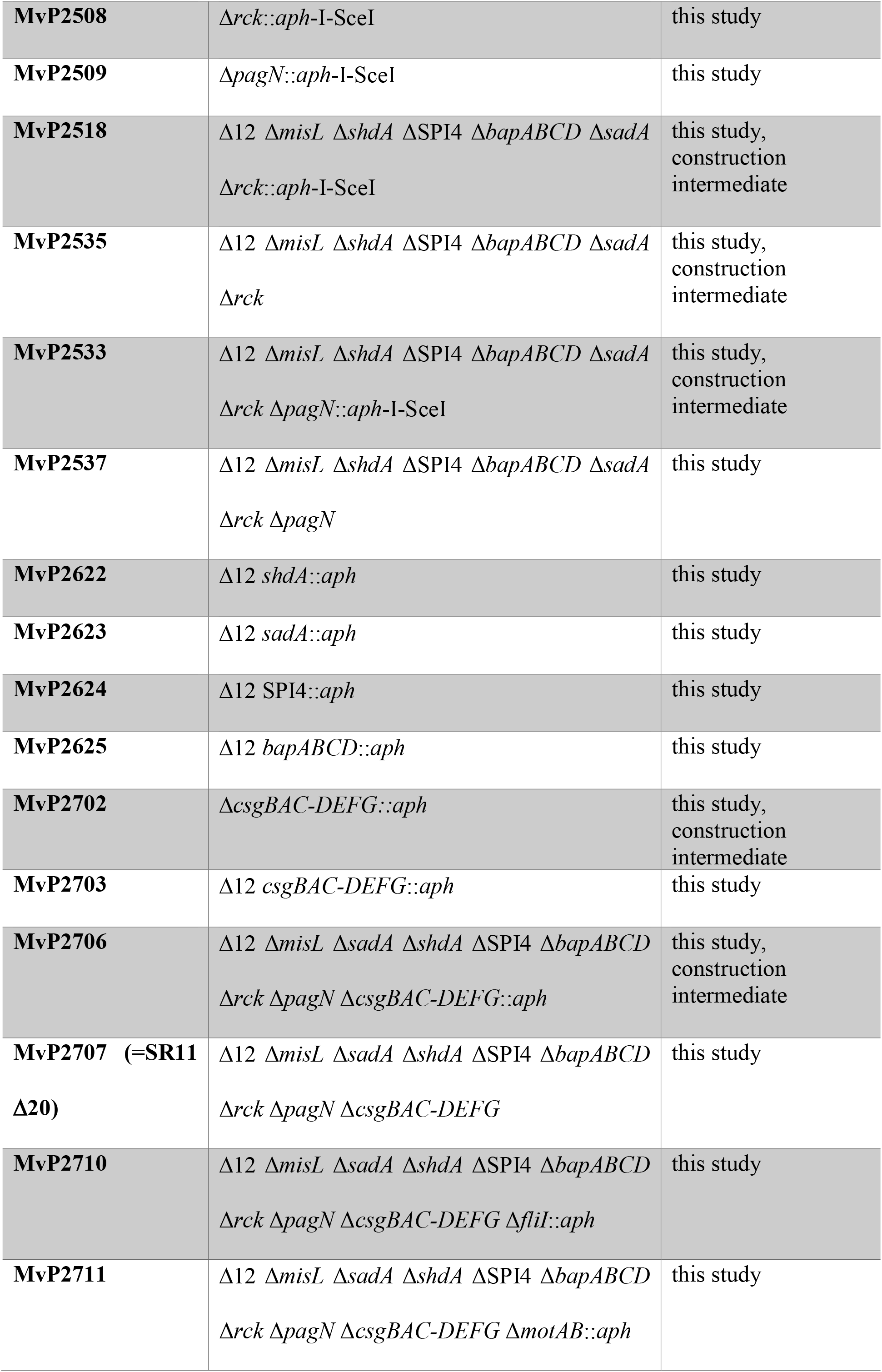
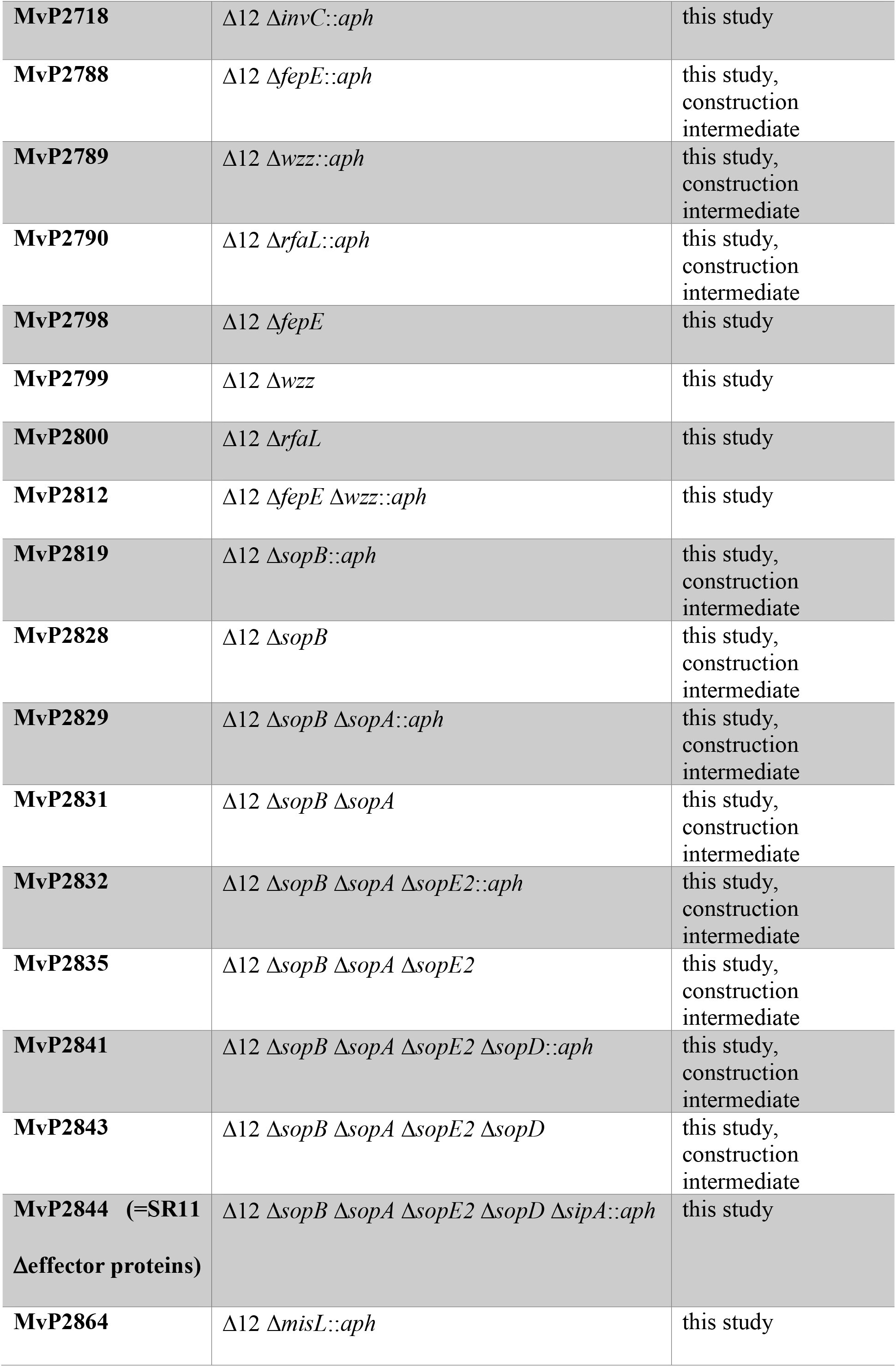
Bacterial strains used in this study.

### Construction of Δ12 strain deleted in chaperone-usher fimbrial gene cluster

Strains are listed in Table 1, plasmids (and the extent of each fimbrial gene cluster deletion) in Table 2, and oligonucleotides in Table S 1. For cloning, *E. coli* DH5α was used as host for pCR2.1 and pBluescriptII-derived plasmids, whereas *E. coli* CC118 λ*pir* was used as host for pRDH10-derived plasmids. To generate the unmarked Δ*lpf*, Δ*pef*, Δ*saf*, Δ*stc*, and Δ*stj* allelic exchange-mediated deletion constructs, an upstream and a downstream region flanking the respective gene cluster to be deleted were amplified from the genome of *S. enterica* serovar Typhimurium LT2 by PCR with primers containing: 1) restriction sites that enable ligation of the flanking regions together at their proximal ends, as well as enable future introduction of an antibiotic resistance cassette; 2) restriction sites to enable subcloning of the deletion construct into the sucrose-counterselectable pRDH10 suicide vector. With the exception of the Δ*lpf* construct, flanking region PCR products were gel purified (QIAEX II Kit; Qiagen), digested with XbaI (NEB), ligated with T4 DNA ligase (NEB), then PCR-amplified by utilizing the distal primer of each respective flanking region’s primer pair. Products were then cloned into pCR2.1 via the TOPO TA kit (Invitrogen), and correct inserts were confirmed by Sanger sequencing (SeqWright). For the *Δlpf* construct, each flanking region was PCR amplified, gel purified, cloned separately into pCR2.1, then confirmed by sequencing. The flanking regions were then joined together by sequential subcloning into pBluescriptII KS+. The unmarked Δ*lpf*, Δ*pef*, Δ*saf*, Δ*stc*, and Δ*stj* constructs were then subcloned into pRDH10. To generate the unmarked Δ*std* and Δ*sti* constructs in pRDH10, the Km-resistance cassette was removed from pEW5 and pEW13, respectively, by restriction digestion, then the vectors were gel purified and religated. As pSF2 (pRDH10 Δ*fim*) did not confer appreciable sucrose sensitivity to strains harboring it, the Δ*fim* construct was subcloned into another site in pRDH10: following *Eco*RI digestion of pSF2, the Δ*fim* construct was gel purified, blunted (QuickBlunt, NEB), and subcloned into the blunted *Bam*HI site of pRDH10, yielding pSPN22. To generate Km-marked deletion constructs, the KSAC cassette of pBS34 was excised with *Xba*I or *Pst*I as relevant, gel purified, then subcloned between the flanking regions of the Δ*lpf*, Δ*pef*, Δ*saf*, Δ*stc*, and Δ*stj* constructs in their respective pRDH10-based vectors. To enable their conjugation, all unmarked and KSAC-marked pRDH10-based fimbrial gene cluster deletion vectors were electroporated into *E. coli* S17-1 λ*pir*.

**Table 2.**
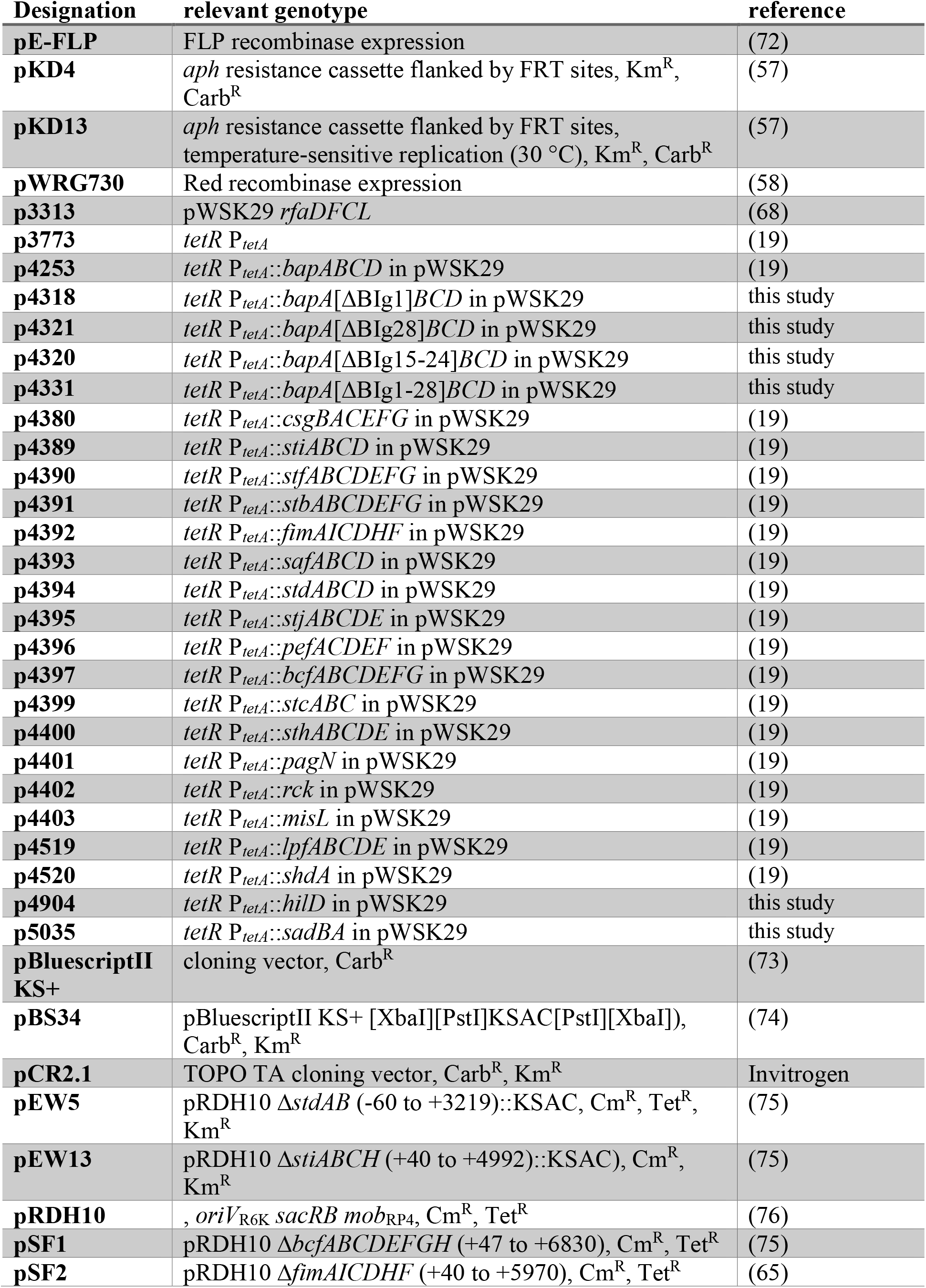
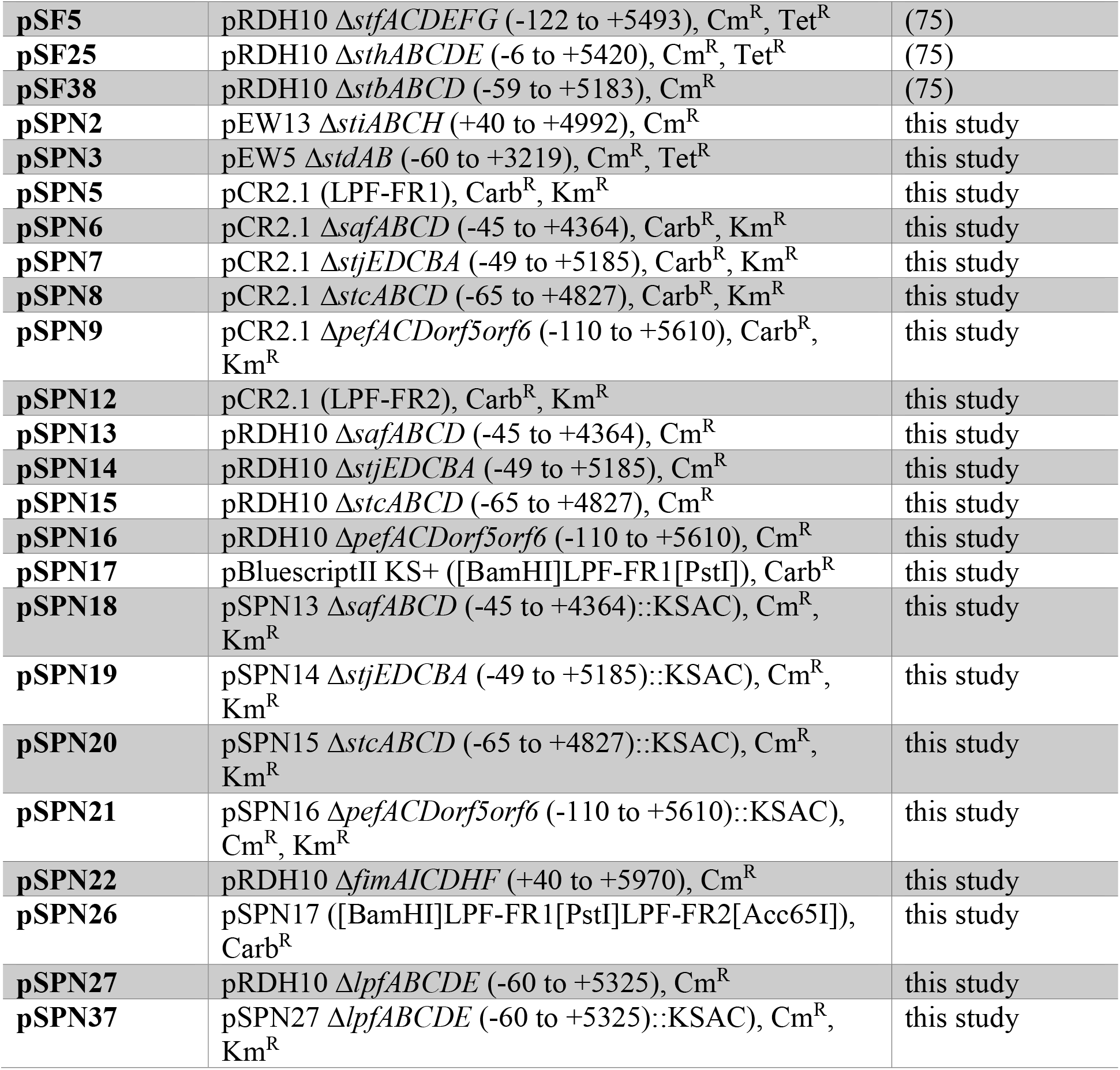
Plasmids used in this study.

*S.* Typhimurium IR715-derived strains harboring a single, KSAC-marked deletion of *lpf*, *pef*, *saf*, *stc*, or *stj* (e.g., SPN195 = IR715 Δ*saf*::KSAC) were generated by conjugation through mating the respective S17-1 λ*pir* pRDH10(Δ::KSAC) strain with IR715. Transconjugants were selected for on LB+Km+Nal agar, and those resulting from a double-crossover event were screened for by sensitivity to Cm, then validated by PCR using primer pairs to confirm that KSAC was located in the correct genomic context, as well as by being negative for PCR amplification of the relevant fimbrial gene cluster’s predicted major subunit gene.

Eleven (*bcf, fim, pef, saf, stb, stc, std, stf, sth, sti, stj*) of the twelve KSAC-marked fimbrial gene cluster deletion strains were then converted to unmarked deletion strains (e.g., SPN230 = IR715 Δ*saf*) by mating the respective S17-1 λ*pir* pRDH10(Δ) and IR715 Δ::KSAC strains. Transconjugants with pRDH10(Δ) integrated into the genome were selected for on LB+Cm+Nal agar, and colonies were then transferred to 5% sucrose agar (55) and incubated at 30 °C. Sucrose-resistant (Suc^R^) colonies lacking the pRDH10(Δ) vector and the Δ::KSAC locus were identified by screening for a Km^S^ Cm^S^ phenotype, and the presence of the unmarked deletion was then validated by obtaining the expected PCR product size when amplifying over the deleted region. To enable transduction of the unmarked deletions (56), we next generated IR715 Δ::pRDH10::Δ strains (e.g., SPN251 = IR715 Δ*saf*::pSPN13), thus reversibly marking the unmarked deletion with the Cm-selectable, sucrose-counterselectable pRDH10 suicide vector. The respective pRDH10(Δ) construct was thus conjugated back into the relevant IR715 unmarked deletion strain, transconjugants with the plasmid integrated into the genome were selected for on LB+Cm+Nal agar, and plasmid integration was further inferred by the inability to PCR amplify across the respective unmarked deletion region due to the size increase.

The *S.* Typhimurium SR11 strain deleted of all 12 chaperone-usher fimbrial gene clusters (Δ12; SPN376) was then generated, with a focus on minimizing the number of passages necessary for introducing each deletion. To begin, Δ*fim*::pSPN22 of SPN227 was transduced via phage P22 HT105/1 *int*-201 into wild-type SR11, and transductants were selected for on LB+Cm agar. As SR11 will accept DNA from P22, but is resistant to lysis by the phage, phage cleanup was unnecessary. Transductants were thus struck immediately to 5% sucrose agar and incubated at 30°C. Suc^R^ colonies were then screened for Cm^S^ by streaking for single colony isolation on both LB and LB+Cm agar. Colony PCR was performed to confirm Δ*fim* status (positive for amplification across the unmarked deletion, negative for *fimA* amplification) of Suc^R^ Cm^S^ colonies. A validated colony was then grown in LB medium, an aliquot of which was used for creating a freezer stock (SPN365 = SR11 Δ*fim*), another aliquot of which was used in the next round of transduction. This process was then repeated for the remaining deletions. The unmarked deletions were transduced first, generating strains SPN366-SPN375. For the final deletion, Δ*lpf*::KSAC of SPN193 was transduced, yielding the Δ12 strain (SPN376). With each successive deletion, every deletion thus far introduced into the strain was reconfirmed by PCR, as was the expected presence/absence of every major fimbrial subunit gene.

### Construction of strains and plasmids

For introduction of the genes *sadBA* under the Tet-on system, template vector p4392 was used harboring *tetR* P*_tetA_*::*fimAICDHF*. Amplification of *sadBA* from the genome of *S*. Typhimurium NCTC 12023 and the vector including the Tet-on system *aph tetR* P*_tetA_* present on p4392 occurred using oligonucleotides as listed in Table S 1 and purified by PCR purification (NEB Monarch). The PCR product encoding for *sadBA* and the PCR product from vector p4392 were assembled by Gibson assembly according to manufactureŕs protocol (NEB Monarch). For overexpression of the *sii* operon, a plasmid was generated for Tet-on expression of transcriptional regulator *hilD*. Using primers listed in Table S 1, *hilD* as amplified from genome *S*. Typhimurium NCTC 12023 genomic DNA, and the vector including *aph tetR* P*_tetA_* present on p4392 was amplified as described before.

Strains with deletion of *csgBAC, csgBAC-DEFG, rck* and *pagN* were created using λ Red recombination in *S*. Typhimurium 12023 harboring pWRG730. One step gene inactivation was performed as described (57) using oligonucleotides as listed in Table S 1. Deletion was checked by colony PCR using oligonucleotides as listed in Table S 1. Further deletion of *aph* was performed using pE-FLP encoding for FLP-recombinase as described (57). For strains lacking *rck* and *pagN* further deletion of *aph* was performed using I-*Sce*I counter-selection as described (58). Generation of strains lacking all fimbrial operons (SR11 Δ12) and one further adhesive structure were created by transferring the deletion by P22 phage transduction. The several deletions were always checked by colony PCR using oligonucleotides as listed in Table S 1.

### Cultivation of sterile grown corn salad

Corn salad seeds (*Valerianella locusta* Verte à cour plein 2, N.L. Chrestensen Erfurter Samen-und Pflanzenzucht) were kindly provided by Dr. Adam Schikora and Dr. Sven Jechalke (Justus Liebig University Giessen). Seeds were sterilized by 70% EtOH for 1 min followed by 3% NaClO for 2 min. Seeds were washed thrice with sterile H_2_O_dd_ and allowed to dry for 30 min. Seeds were planted on Murashige-Skoog (MS-) agar (per liter: 1.1 g Murashige-Skoog medium including vitamins, Duchefa Biochemie #M0222; 1 g agar; 0.5 g MES; pH 5.4) in sterile plastic containers with air filter (round model 140mm; Duchefa Biochemie #E1674) at 20 °C with a 12 h/12 h day/night-rhythm for 8 weeks.

### Adhesion to corn salad

For infection of corn salad by *Salmonella* leaf discs of 8 mm average of 8 weeks old plants were punched out by biopsy punches immediately before infection process. 48-well plates were used with one leaf disc per well mechanically fixed by sterile stainless-steel inlays. For each condition, three leaf discs were infected. For infection o/n cultures of *Salmonella* strains were diluted 1:31 in LB (containing antibiotics if required) and grown for 3.5 h in test tubes with aeration in a roller drum. The cultures were diluted in PBS to obtain approximately 5.6 x 10^7^ bacteria/ml and 50 µl of this inoculum was spotted onto one leaf disc. Infection process occurred either for 1 h, RT under static conditions or for 55 min, RT after a centrifugation step 500 x g for 5 min. After infection, leaf discs were washed once with PBS to remove non-bound bacteria. Three leaf discs were transferred to tubes and washed two further times with PBS by short mixing on a Vortex mixer. Plant tissue was homogenized with a pellet pestle motor in 600 µl 1% sodium deoxycholate in PBS and colony forming units were determined by plating serial dilutions of the lysates on MH agar plates (Müller-Hinton agar plates) incubated o/n at 37 °C. A non-infected sample was used in every assay to ensure the sterility of the corn salad.

### Flow cytometry

For analysis of surface expression of SadA and BapA by flow cytometry 6×10^8^ bacteria were washed in PBS and then fixed with 3% paraformaldehyde/PBS for 20 min. Bacteria were blocked with 2% goat serum in PBS for 30 min and afterwards stained with the specific antiserum goat-α−SadA or goat-α-BapA diluted 1:250 and 1:1,000 in 2% goat serum/PBS for 2 h and goat α rabbit IgG antibody coupled to Alexa-Fluor488 diluted 1:2,000 in 2% goat serum/PBS for 1 h. For analysis of surface expression of SiiE by flow cytometry, ca. 3×10^8^ bacteria were fixed in 3% paraformaldehyde in PBS for 20 min. Bacteria were blocked with blocking solution (2% goat serum and 2% bovine serum albumin in PBS) for 30 min and afterwards stained with the specific antiserum α-SiiE-C-terminal coupled to Alexa-Fluor488 (1:100) for 1 h. Bacteria were measured with a Attune NxT Flow Cytometer (Thermo Fisher) and analyzed using Attune NxT Software version 2.7. A mutant strain lacking the respective adhesive structure was used as a negative control for gating.

## Acknowledgements

This work was supported by the Bundesanstalt für Landwirtschaft und Ernährung (BLE) by project *plantinfect,* grant 2813HS027). Further support by the DFG by grant SFB 944, project Z is kindly acknowledged. We thank the members of the *plantinfect* consortium for fruitful discussion and exchange of reagents. We like to thank Inigo Lasa and Dirk Linke for sharing antisera against BapA and SadA, respectively.

## Suppl. Tables

Table S 1. Oligonucleotides used in this study

## Suppl. Figure Legends

**Figure S 1: Schematic overview of adhesion assay for the infection of corn salad.** Sterile plant tissue (A). Punching out several leaf discs using a biopsy punch with a diameter of 8 mm. Leaf discs were used immediately. (B). Each leaf disc (28.3 mm^2^) was infected with 2.81 x 10^5^ bacteria for 1 h at RT (C), under static conditions or with forced contact by centrifugation at 500 x g for 5 min (D). Removal of non-bound bacteria by washing once with PBS (E). Transfer of plant tissue in tubes (F). Removal of non-bound bacteria by washing twice with PBS and short mixing on a Vortex mixer (G). Homogenization of leaf discs in 1 % deoxycholate/PBS using a pellet pestle motor (H). Plating homogenates and the inoculum onto agar plates for colony growth (I). Quantification of adhesion rates (J).

**Figure S 2: Quantification of surface expression of SiiE, BapA and SadA by flow cytometry.** Tet-on expression of adhesins SiiE (A, B), BapA (B, C), and SadA (E, F) was measured by flow cytometry. Adhesins were detected using antisera rabbit α-SiiE (1:1,000), rabbit α-BapA (1:1,000), or rabbit α-SadA (1:250). As secondary antibody, rabbit α-goat-Alexa488 (1:2,000) was used. In A), C), and E), overlays of the measured fluorescence intensities are shown, whereas in B), D), and F), the percentages of Alexa488-positive bacteria are shown.

**Figure S 3: Microscopic analysis of SR11 Δ12 expressing WT BapA and truncated forms of BapA.** 3.5 h subcultures induced with AHT or not induced were diluted to 1 x 10^8^ bacteria/ml in PBS. Bacteria were visualized using a Zeiss Axio Observer with brightfield microscopy with a 40x objective. Images were recorded with a AxioCam and data were process in ZEN 2012. Scale bars, 20 µm.

